# Inhibition of dipeptidyl peptidase 4 allows accurate measurement of GLP-1 secretion in mice

**DOI:** 10.1101/2023.10.19.563029

**Authors:** Mark M. Smits, Katrine D. Galsgaard, Sara Lind Jepsen, Nicolai Wewer Albrechtsen, Bolette Hartmann, Jens J. Holst

## Abstract

Dipeptidyl peptidase (DPP)-4 and neprilysin (NEP) within few minutes degrade glucagon-like peptide 1 (GLP-1) in mice, generating small, inactive fragments of the peptide. Commercially available sandwich ELISA kits may not accurately detect intact GLP-1 (that is GLP-1[7–36]NH_2_) and these moieties, leading to underestimation of secretion and potentially misleading results. Single-site antibody approaches may pick up some fragments, yet require large plasma volumes and the accuracy is uncertain.

We aimed to find a way to stabilize GLP-1 in mice allowing reliable measurement with sensitive commercially available ELISA kits. Non-anesthetized male C57Bl/6JRj mice were subjected to an oral glucose tolerance test (OGTT; 2 g/kg glucose via oral gavage). Blood was drawn from the retrobulbar plexus before and repeatedly during the OGTT and total and intact GLP-1 were measured by commercially available sandwich ELISA kits (Mercodia and Alpco, respectively). In blood samples taken 15 minutes after the glucose load, there were no increase in plasma GLP-1 concentration. There was a small insignificant increase in total GLP-1 (1-2 pmol/L) in samples taken at t=5 and t=10 minutes following the OGTT, but no rise in intact GLP-1. We then administered control (saline), or a DPP-4 inhibitor (valine pyrrolidide, 0.1 µmol/g or sitagliptin, 10 mg/kg) with or without a NEP-inhibitor (sacubitril, 0.3 mg/kg) 30 minutes before the OGTT. In the four inhibitor groups, intact GLP-1 increased during the OGTT (levels ranging between 4.04 [SD 2.87] and 15.53 [SD 6.76] pmol/L). The combination of sitagliptin with sacubitril gave the largest increase in intact GLP-1 levels. Finally, after injecting male C57Bl/6JRj mice with a known dose of GLP-1(7–36)NH_2_, the peak GLP-1 levels were significantly higher during sitagliptin, but not with the combination of sitagliptin/sacubitril. Both inhibitor groups, however, showed prolonged half-life of the GLP-1 plasma disappearance.

We conclude that for measurements of GLP-1 secretion in mice with commercially available sandwich ELISA kits, it is necessary to consider both timing of blood sampling and *in vivo* inhibition of DPP-4. The described approach allows improved estimates of GLP-1 secretion for future studies. It is a limitation that DPP-4 and NEP inhibition may have metabolic effects by stabilizing the intact GLP-1 peptide (e.g. influencing levels of insulin and glucagon).

## Introduction

The gut hormone glucagon-like peptide-1 (GLP-1) is secreted from enteroendocrine L-cells of the gastrointestinal mucosa in response to a variety of stimuli (1). It is secreted in its active, or intact form, GLP-1(7–36)NH_2_, however, as degradation starts immediately after secretion and at a high rate, it is estimated that only 7-8% of the intact GLP-1 reaches the systemic circulation (2). Degradation occurs at the N-terminus, catalysed by the ubiquitous enzyme dipeptidyl peptidase-4 (DPP-4) which is present in intestinal capillaries, resulting in formation of the generally inactive metabolite GLP-1(9–36)NH_2_. In addition, in mice there is pronounced additional endoproteolytic degradation by neprilysin (neutral endopeptidase 24.11 or NEP), which strongly reduces the stability of the inactive metabolite GLP-1(9–36)NH_2_ (***Supplemental Figure 1a***) (3).

The degradation pattern of GLP-1 influences the ways by which it can be measured. The most frequently used method to measure GLP-1 is by immunoassays. Both radioimmunoassays (RIA) and the various enzyme-linked immunosorbent assays (ELISA) rely on antibodies against epitopes of GLP-1. The binding sites of these antibodies may be directed against the terminals (‘terminal wrapping antibodies’), potentially allowing differentiation between the different molecular forms. Hence, terminal-wrapping antibodies against the N-terminus might distinguish between intact GLP-1(7–36)NH_2_ and the primary metabolite of DPP-4 mediated degradation, GLP-1(9–36)NH_2_. Antibodies against the mid-region would be expected to measure all GLP-1 forms (including larger incompletely processed forms, as well as active and inactive forms), whereas antibodies directed against the C-terminus would measure all forms of which this region is exposed by enzymatic processing. In humans and mice, most GLP-1 is amidated at the C-terminus (4), which is why terminal-wrapping antibodies against the amidated C-terminus allow specific determination of molecules with this modification in these species. In other species (rats and pigs), amidation is incomplete (4), complicating assays based on C-terminal antibodies. Thus, designing assays for reliable GLP-1 measurements is by no means simple.

Importantly, many of the commercially available kits are based on the ‘sandwich’ technique, which uses antibodies against two different epitopes of GLP-1, such as antibodies directed against each of the termini. This may work in humans, provided that the antibodies fully recognize GLP-1(9–36)NH_2_, which is relatively stable in humans with a plasma half-life of about 5 min. However, this technique frequently seems to fail in mice, where this metabolite is further fragmented by NEP (3) (***Supplemental Figure 1b***). As a result, many investigators experience situations where otherwise reliable stimuli do not result in the expected significant GLP-1 increases (as seen in (3, 5)). Moreover, in those cases where GLP-1 levels do increase after stimulation, the absolute levels are below what could be expected (for example in (6)). While this problem could be circumvented by using single-site techniques for one of these fragments, assuming that these fragments are adequately immunoreactive, these assays typically need large plasma volumes (at least 100 µL), allowing only a single measurement per mouse (***Supplemental Figure 1c***). Alternatively, measuring NEP-created fragments, like GLP-1(28–36)NH_2_, could be useful, but sensitive sandwich-assays for this are not currently available.

In the present study, we investigated several approaches to estimate plasma GLP-1 levels more accurately in mice with commercially available kits.

## Methods

### Animal Studies

All studies were conducted with permission from the Danish Animal Experiments Inspectorate (2018-15-0201-01397) and the local ethical committee in accordance with the guidelines of Danish legislation governing animal experimentation (1987) and the National Institute of Health. Male C57Bl6/JRj mice (Janvier labs), aged 12-17 weeks, were housed in groups of 4 to 6 in individually ventilated cages with a 12 h light cycle (lights on 06:00–18:00) and ad libitum access to standard chow and water. Experiments were performed after at least 1 week of acclimatization.

### Oral glucose tolerance tests

Several studies were performed based on an OGTT, following the same protocol; After a 6 hour fast (08:00-14:00), D-glucose at 2 g/kg body weight (50% wt/vol; Amgros I/S; Copenhagen) was given to non-anesthetized mice through oral gavage. Blood glucose was measured directly from tail tip puncture just before the oral glucose load, and repeatedly thereafter using a handheld glucometer (Accu-Chek Mobile U1; F. Hoffmann–La Roche, Basel, Switzerland). Time-points varied per experiment.

To see whether ex vivo stabilization of GLP-1 is sufficient to measure the hormone, blood was drawn from the retro-bulbar plexus (∼75 µL), using EDTA-coated capillary tubes, immediately after the blood glucose measurements, and immediately transferred to pre-chilled plastic tubes containing inhibitor-cocktail, which was then vortexed briefly to ensure proper mixing. The cocktail consisted of the DPP-4 inhibitor valine pyrrolidide (gift from Novo Nordisk, Denmark) and the serine protease inhibitor Pefabloc (Thermo Scientific), both dissolved in Milli-Q water, and reaching a final concentration of 0.01 mmol/L and 1 mg/mL, respectively. Tubes were kept on ice until centrifugation, after which plasma was transferred to PCR tubes and stored at -20°C until analysis.

To see whether *in vivo* inhibition of GLP-1 degrading enzymes leads to measurable GLP-1 levels, enzyme inhibitors or control (PBS) were given 30 minutes before administration of glucose. The NEP-inhibitor sacubitril (AHU-377; Nordic BioSite, Denmark) was dissolved in DMSO to a concentration of 10 mg/mL as stock solution. On experimental days, a 0.14 mmol/L working solution was made in phosphate-buffered saline (PBS), to allow oral gavage at 0.3 mg/kg body weight. The DPP-4 inhibitor valine pyrrolidide was dissolved in PBS to a concentration of 4.12 mg/mL for oral administration (20.6 mg/kg body weight) and 5 mg/mL for i.p. administration (15 mg/kg). Sitagliptin (Xelevia; Merck, Sharp & Dohme) was dissolved in PBS to a concentration of 2 mg/mL, allowing oral administration at 10 mg/kg body weight. These inhibitors were tested in several combinations, all given at a (oral) dose of 10 µL/g body weight. Since the final concentration of DMSO in the sacubitril-containing combinations was 0.3%, we added DMSO in that concentration to all solutions without sacubitril (i.e. controls and DPP-4 inhibitor only) to balance this factor.

In one set of experiments, we added acetaminophen (0.1 mg/g) to the glucose solution, to assess gastric emptying rate.

### *In vivo* disappearance of GLP-1(7–36)NH_2_

In two separate experiments, we tested the *in vivo* disappearance of exogenously administered GLP-1(7–36)NH_2_, and addressed whether *in vivo* enzyme inhibition could decrease disappearance using the inhibitors as outlined above, administered 30 minutes before the injection of GLP-1(7–36)NH_2_. A stock solution of 50 µg/mL, dissolved in 1% albumin in MilliQ-water, was made with synthetic GLP-1(7–36)NH_2_ (Bachem, Switzerland). On the day of the experiment, the stock solution was diluted with PBS, allowing administration of 800 fmol in 100 µL. In the first experiment, GLP-1(7–36)NH_2_ was injected into a tail vein of non-anesthetized mice. Blood was drawn from the retro-bulbar plexus just before the injection and 1, 2, 3 and 10 minutes following the injection. In the second experiment, animals were sedated with isoflurane (induction 5%, maintenance 2.5%), and after laparotomy, GLP-1(7–36)NH_2_ was injected into the vena cava. Blood was drawn from the retro-bulbar plexus 2, 4, 6 and 10 minutes following the injection.

### *In vitro* plasma disappearance of GLP-1(7–36)NH_2_ and GLP-1(9–36)NH_2_

We assessed the *in vitro* plasma disappearance of GLP-1(7–36)NH_2_ and GLP-1(9–36)NH_2_ (Bachem, Switzerland) independent of use of inhibitors. Six mice were sedated using isoflurane, and total blood volume was drawn from the vena cava into EDTA-coated tubes. After centrifuging, plasma was immediately separated and chilled. Just before the experiment, plasma was pooled and distributed between 4 tubes (600 µL per tube), and warmed to 37°C. Synthetic GLP-1 was added to each plasma tube (final concentration 200 pmol/L), after which 60 uL samples were withdrawn after 0, 1, 2, 5 10, 30, 60, 120 and 180 minutes incubation. Samples were transferred to prechilled tubes containing inhibitors (valine pyrrolidide and Pefabloc as described above), after which they were snap frozen in liquid nitrogen.

### Biochemical measurements

We used several sandwich ELISA kits to measure total and intact GLP-1. For total GLP-1 measurements, involving side-viewing antibodies binding near the termini, we used MSD (cat.nr. K1503PD; RRID: AB_2814818) and Mercodia (cat.nr. 10-1278-01; RRID: AB_2892202) kits. For intact GLP-1 measurements, where a side-viewing antibody binds the N-terminus (and the other antibody near the C-terminus) we used MSD (cat.nr. K1503OD; RRID: AB_2935695) and Alpco (cat. nr. 80-GLP1A-CH01; RRID: AB_2941993) kits. Finally, in one experiment we measured GLP-1[9-36]NH_2_ using a kit from JBL (cat.nr. JP27788; RRID: AB_3065262).

Mercodia ELISA kits were used for measurement of insulin (cat.nr. 10-1247-01; RRID: AB_2783837) and glucagon (cat.nr. 10-1281-01; RRID: AB_2783839). Acetaminophen was measured using a spectrophotometric assay (Sekisui, cat.nr. Acetaminophen-L3K). All assays were run according to the manufacturer’s instructions.

### Statistical analysis

Our primary endpoint was to assess whether any of the interventions could lead to a measurable increase in GLP-1 levels coinciding with an increase in glucose and insulin. Statistical analyses were performed using Graphpad Prism v9.5 (Graphpad software LLC, San Diego, USA). Data are shown as mean ± standard deviation (SD) to describe values, whereas comparisons between groups are shown as mean (95% confidence interval). To compare values between two groups at a single time-point, a t-test was used. When comparing more than one group, a one-way ANOVA with post-hoc Šidák was employed. To test an increase of GLP-1 compared to baseline, a one-way repeated measures ANOVA with post-hoc Šidák was used. A two-sided p-value <0.05 was considered statistically significant.

## Results

### For total GLP-1, timing might be key

We first performed an OGTT and measured total GLP-1 using the Mercodia kit in plasma samples taken immediately before the glucose load (0 minutes) and 15, 30 and 60 minutes after. Despite an adequate increase in plasma glucose levels (***Figure 1a***), we did not observe an increase in total GLP-1 levels after glucose stimulation. Moreover, the GLP-1 levels were generally low (<0.9 pmol/L; where the detection limit of the Mercodia-kit is 0.65 pmol/L) (***Figure 1b***). Before including enzyme inhibitors, we tried different sampling time-points, as the total GLP-1 peak after an OGTT might actually occur before 15 minutes. Indeed, a statistical significant increase in total GLP-1 could be found at 5 minutes (and a numerical increase at 10 minutes) (***Figure 1c***). However, the absolute increases were low (∼ 3 pmol/L).

**Figure 1.**
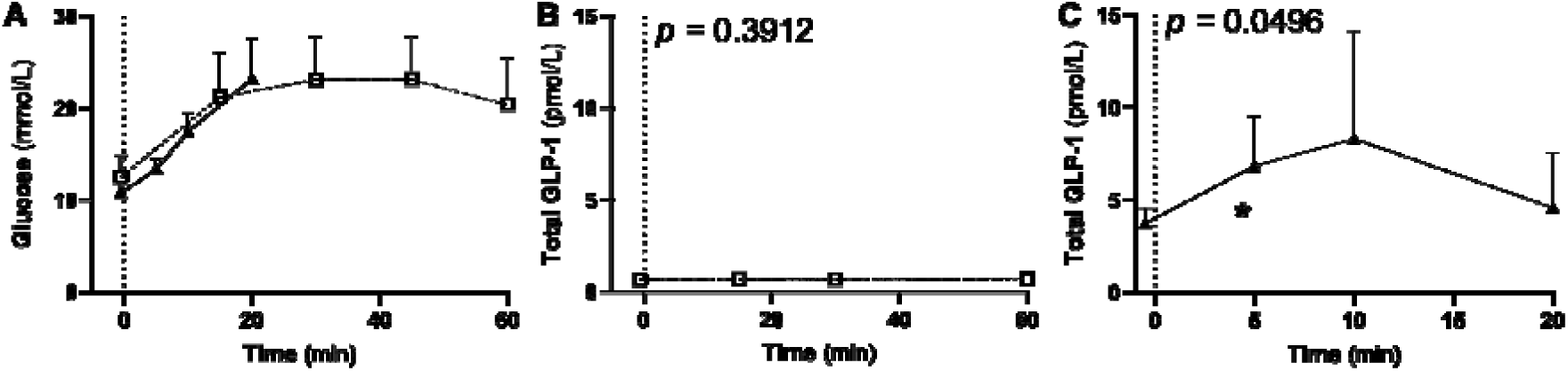
Timing of blood withdrawal might be crucial for measuring total GLP-1. **Legend:** In two sets of experiments, total GLP-1 levels were measured using the Mercodia ELISA kit (cat.nr. 10-1278-01) before and after an oral glucose load (given at t=0 min) in male C57Bl/6JRj mice. *Panel (A)* shows that the OGTT increases glucose levels. Since GLP-1 secretion occurs in response to glucose absorption, GLP-1 should increase simultaneously. The dashed line corresponds to the study as shown in panel B, the solid line to panel C. *Panel (B)* shows the total GLP-1 levels at t=0, 15, 30 and 60 minutes (n=7). The glucose curve for this experiment is shown in A with a similar dashed line. In *panel (C)* GLP-1 levels are shown at 5, 10 and 20 minutes after an OGTT (n=8). Values are shown as mean ± SD. Statistics were performed using one-way repeated-measures ANOVA. Overall p-values are given, and an asterisk indicates a difference from baseline at that time point (\**p*<0.05)

We then used the optimized time-points to measure intact GLP-1, using the Alpco kit (***Figure 2a***). Despite the optimized sampling time-points, intact GLP-1 levels remained low (<4.6 pmol/L), and no significant increase was seen after stimulation by glucose (***Figure 2b***).

**Figure 2.**
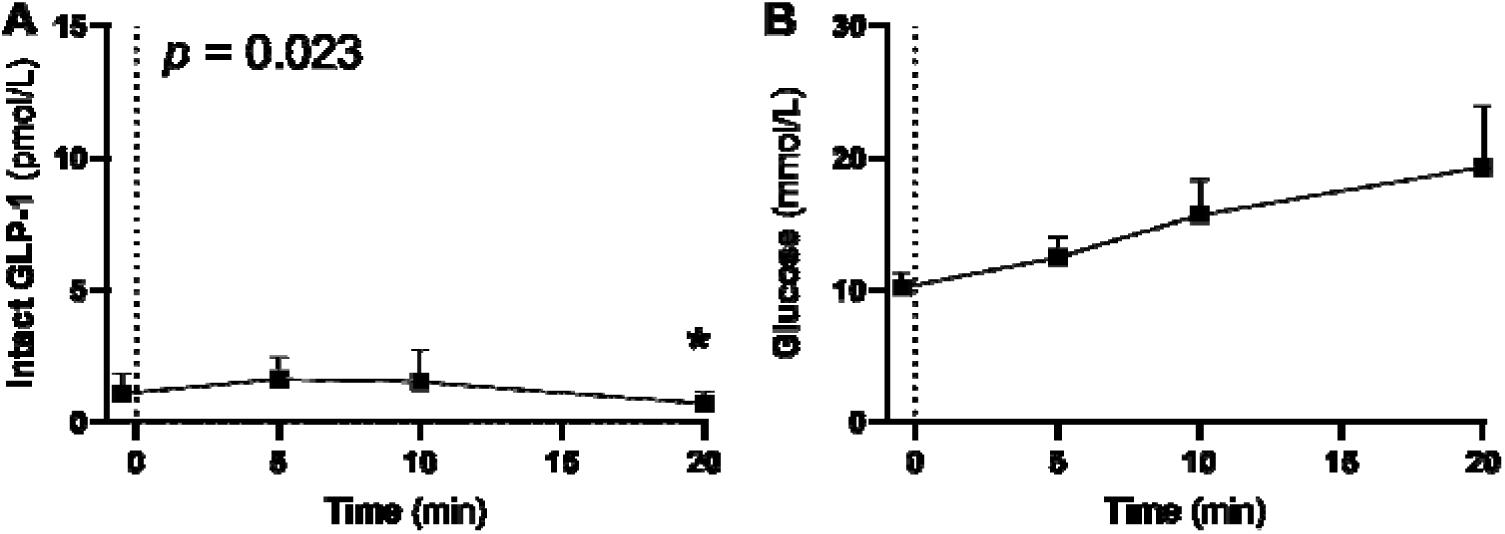
Early blood sampling is not sufficient for intact GLP-1 measurement. **Legend:** An OGTT was performed in male C57Bl/6JRj mice (n=12), and blood was taken at time t=0, 5, 10 and 20 min. *Panel (A)* shows the intact GLP-1 levels measured by the Alpco kit (Cat.nr. 80-GLP1A-CH01). Values are shown as mean ± SD. Statistics were performed using one-way repeated-measures ANOVA. Overall p-values are given, and an asterisk indicates a difference from baseline at that time point (\**p*<0.05). *Panel (B)* shows the glucose levels.

### Ex vivo enzyme inhibition is not sufficient to measure intact GLP-1

We then set out to see whether inhibition of GLP-1 degrading enzymes in the blood collection tube could work as previously described (7). Adding a combination of valine pyrrolidide and Pefabloc to *the collection tube* yielded very similar results compared to when no inhibitors were added (the control group) (***Figure 3a*** and ***3b***). Glucose levels increased similarly in both groups (***Figure 3c***).

**Figure 3.**
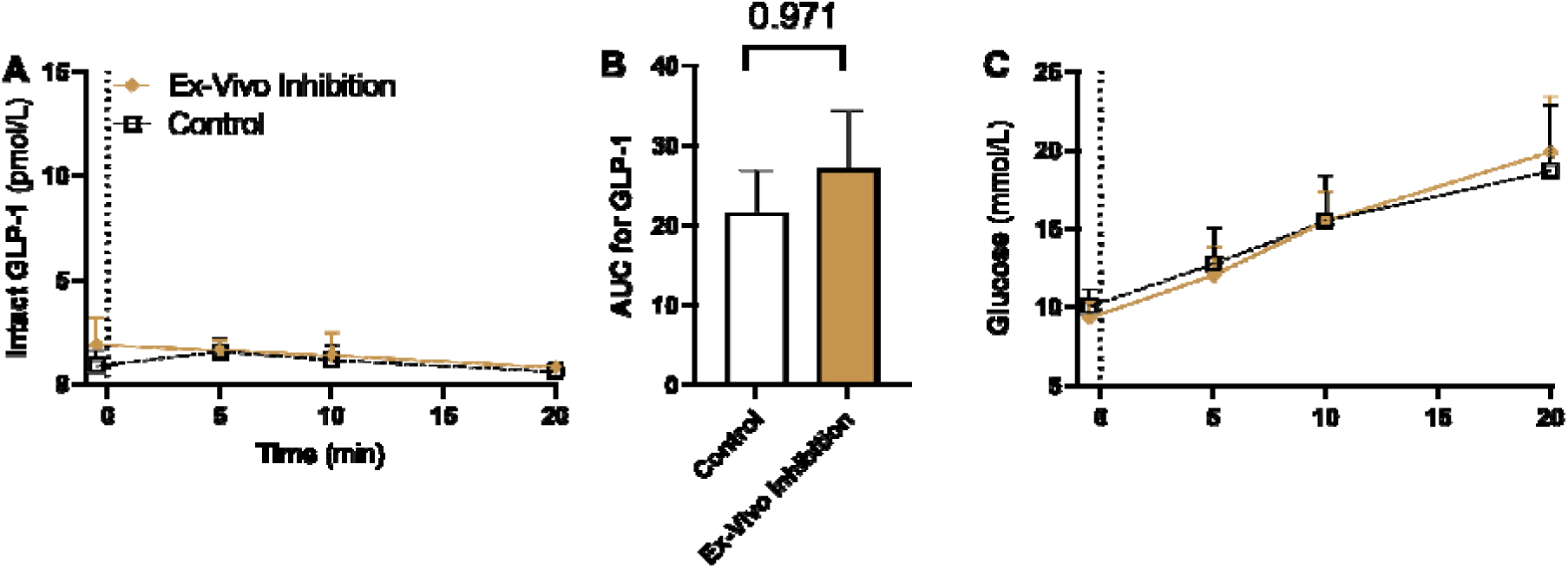
Ex vivo inhibition is not sufficient to measure intact GLP-1 in mice. **Legend:** Effects of ex vivo enzyme inhibition using the DPP-4 inhibitor Valine pyrrolidide and protease inhibitor pefabloc (brown lines; n=6) and no inhibition (control; black dashed line; n=9) after an oral glucose load given at t=0 min in male C57Bl/6JRj mice. *Panel (A)* shows intact GLP-1 as measured by the Alpco kit, and in *Panel (B)* the total area under the curve for intact GLP-1. No differences were observed between the two groups; *Panel (C)* shows the corresponding glucose levels taken from the tail tip, a different sample than the one used for GLP-1 measurements, and thus not subjected to enzyme inhibitors. Values are shown as mean ± SD. Statistics were performed for the AUC using an unpaired t-test.

### *In vivo* inhibition of DPP-4 and NEP gives a measurable intact GLP-1 response to an oral glucose load

Next, we tested our hypothesis that degradation of intact GLP-1 occurs so rapidly, that *in vivo* enzyme inhibition is needed to get measurable levels.

First, we tested the NEP inhibitor sacubitril alone, given orally 30 minutes before the OGTT (***Figure 4a***). GLP-1 levels were not different compared to the control group (difference in AUC of 4.0 [-65.7 to 72.8] pmol/L/min; p>0.99).

**Figure 4.**
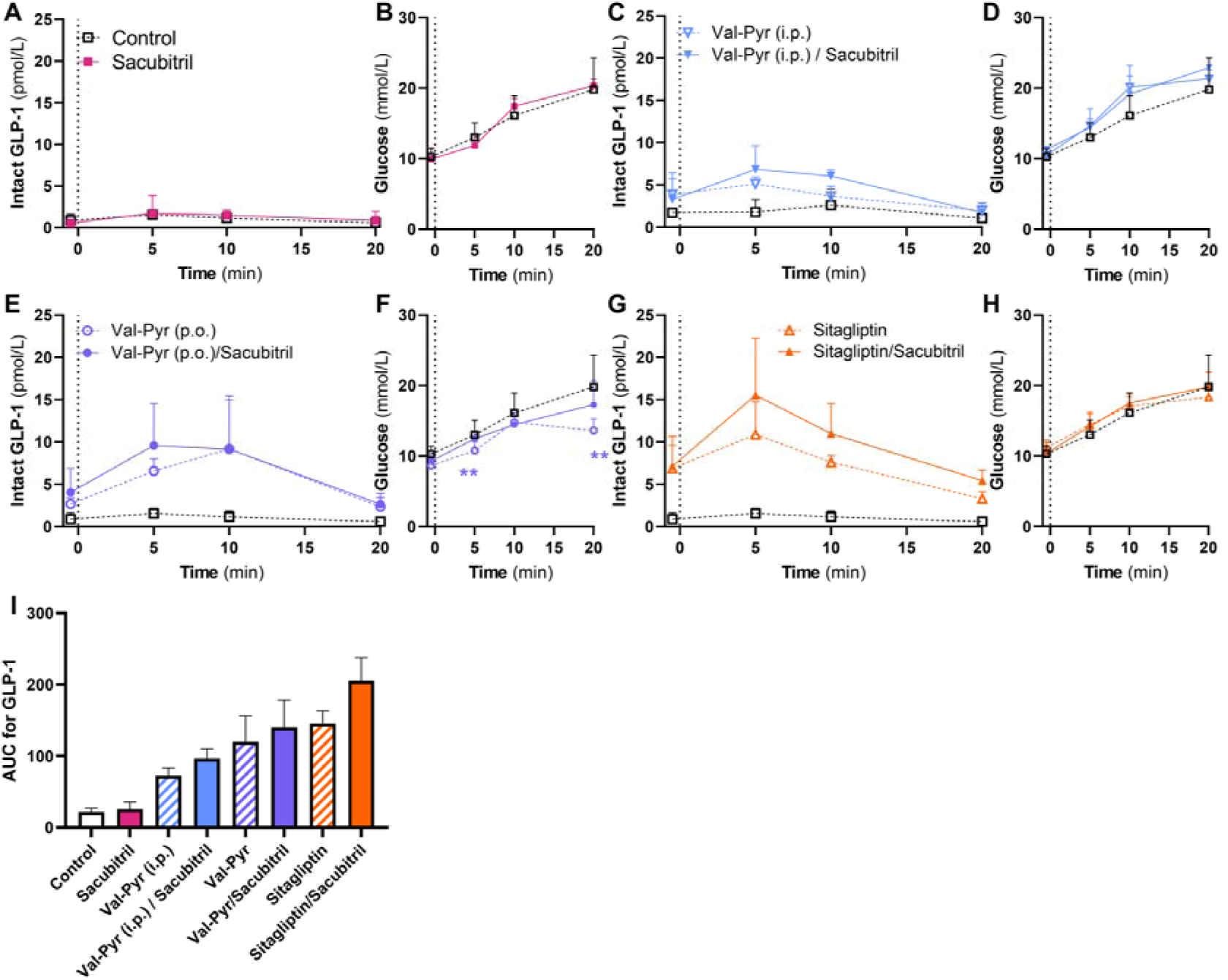
DPP-4 needs to be inhibited *in vivo* to measure intact GLP-1 in mice. **Legend:** Effects of *in vivo* inhibition of DPP-4 and/or NEP on levels of intact GLP-1 (as measured with the Alpco kit) after an OGTT in male C57Bl/6JRj mice. Oral glucose was given at t=0, and blood was drawn from the retrobulbar plexus at t = 0-, 5-, 10- and 20-min. Different combinations of DPP-4 inhibitors, with or without sacubitril (NEP inhibitor), are shown. *Panel (A)*: Intact GLP-1 levels during control situation (no inhibitors, dashed black line) and during NEP inhibition only (sacubitril, magenta solid line; closed square, n=6). *Panel (B)*: the glucose curve for panel A; *Panel (C)* shows intact GLP-1 levels after DPP-4 inhibitor Valine pyrrolidide given intraperitoneally (i.p.) (blue dashed line, open triangle, n=5), the combination of Valine pyrrolidide i.p. with oral sacubitril (blue line, closed triangle, n=5) or control (black dashed line, open square); *Panel (D)*: the glucose curve for panel C; *Panel (E)* shows intact GLP-1 levels after Valine pyrrolidide given orally (p.o.) (purple dashed line, open circle, n=5), or the combination of oral Valine pyrrolidide with sacubitril (purple line, closed circle, n=5); *Panel (F)*: the glucose curve for panel E; *Panel (G)* shows intact GLP-1 levels after DPP-4 inhibitor sitagliptin given orally (orange dashed line, open triangle, n=6), or the combination of sitagliptin with sacubitril (orange line, closed triangle, n=6); *Panel (H)*: the glucose curve for panel G; *Panel (I)* shows the total area under the curves of intact GLP-1 based on panels A-D; Values are shown as mean ± SD. Statistical analyses were performed for the AUCs in panel D, using one-way ANOVA with post-hoc Šidák focusing only on the difference between inhibitors and control, and the inhibitor-combinations (DPP-4 and NEP inhibitor) versus the respective DPP4 inhibitor only. Analyses in panel F were performed using two-way ANOVA with post-hoc Šidák, asterisks indicate a difference from control at that time point (\*\**p*<0.01).

In contrast, when giving a DPP-4 inhibitor orally 30 minutes before the OGTT, levels of intact GLP-1 increased upon glucose stimulation (***Figures 4b-d***). With valine pyrrolidide given intraperitoneally (***Figure 4b***) or orally (***Figure 4c***), fasting intact GLP-1 levels doubled compared to controls, yet the absolute levels were still low (from to 1.7 to 3.9 pmol/L). After the OGTT, levels rose to 9.2±6.3 pmol/L (at 10 minutes) with oral valine pyrrolidide (***Figure 4c***), while i.p. valine pyrrolidide retained modest levels of 5.1±0.9 pmol/L (***Figure 4b***). With sitagliptin (***Figure 4d***), both fasting (6.9±3.8 pmol/L) and post-OGTT (10.9±3.9 pmol/L at 5 minutes) intact GLP-1 levels were increased. However, the largest increases in intact GLP-1 were obtained by combining a DPP-4 inhibitor and sacubitril, particularly the combination with sitagliptin (***Figure 4b-d***). Sitagliptin/sacubitril treatment gave the highest total AUC of intact GLP-1 of the tested combinations (an increase of 850% [536 to 1169] versus control; p<0.0001) (***Figure 4e***). Importantly, during all tested conditions, glucose levels rose (***Figure 4f***), and thus an increase in intact GLP-1 was expected. It must be noted, however, that the groups with DPP-4 inhibitors had a suppression in the rise of glucose at 20 minutes, which was statistically significant for valine pyrrolidide only (−6.2 [-1.9 to -10.4] mmol/L, p=0.0023).

We then repeated some of the *in vivo* inhibition studies and measured total GLP-1 using the Mercodia kit, to see whether this would result in increased GLP-1 levels (***Supplemental figure 2***). However, neither inhibition of NEP alone, DPP-4 alone, nor the combination of both, increased GLP-1 levels compared to the control situation.

### *In vivo* DDP-4 and NEP inhibition may affect blood glucose, glucagon levels and gastric emptying during the OGTT

Inhibition of DPP-4 and NEP affects the cleavage metabolism of several hormones, including GLP-1 and glucagon (8–11). For example, DPP-4 inhibition may reduce glucagon levels, while NEP-inhibition increases them (8–11). As such, *in vivo* enzyme inhibition with the aim of measuring intact GLP-1 may affect other physiological parameters which are normally measured during an OGTT. Therefore, in one set of experiments with oral valine pyrrolidide and/or sacubitril, we additionally measured plasma levels of insulin, glucagon and acetaminophen (***Figure 5***). Although total AUCs for glucose (***Figure 5a***), insulin (***Figure 5b***) and glucagon (***Figure 5c***) were not statistically different between inhibitor-groups and control, they were numerically different. We added acetaminophen to the glucose load to estimate potential differences in gastric emptying rate (***Figure 5d***). While total AUCs were similar between groups, the levels seemed to rise slightly faster in the combinations with sacubitril, indicating an increased gastric emptying rate.

**Figure 5.**
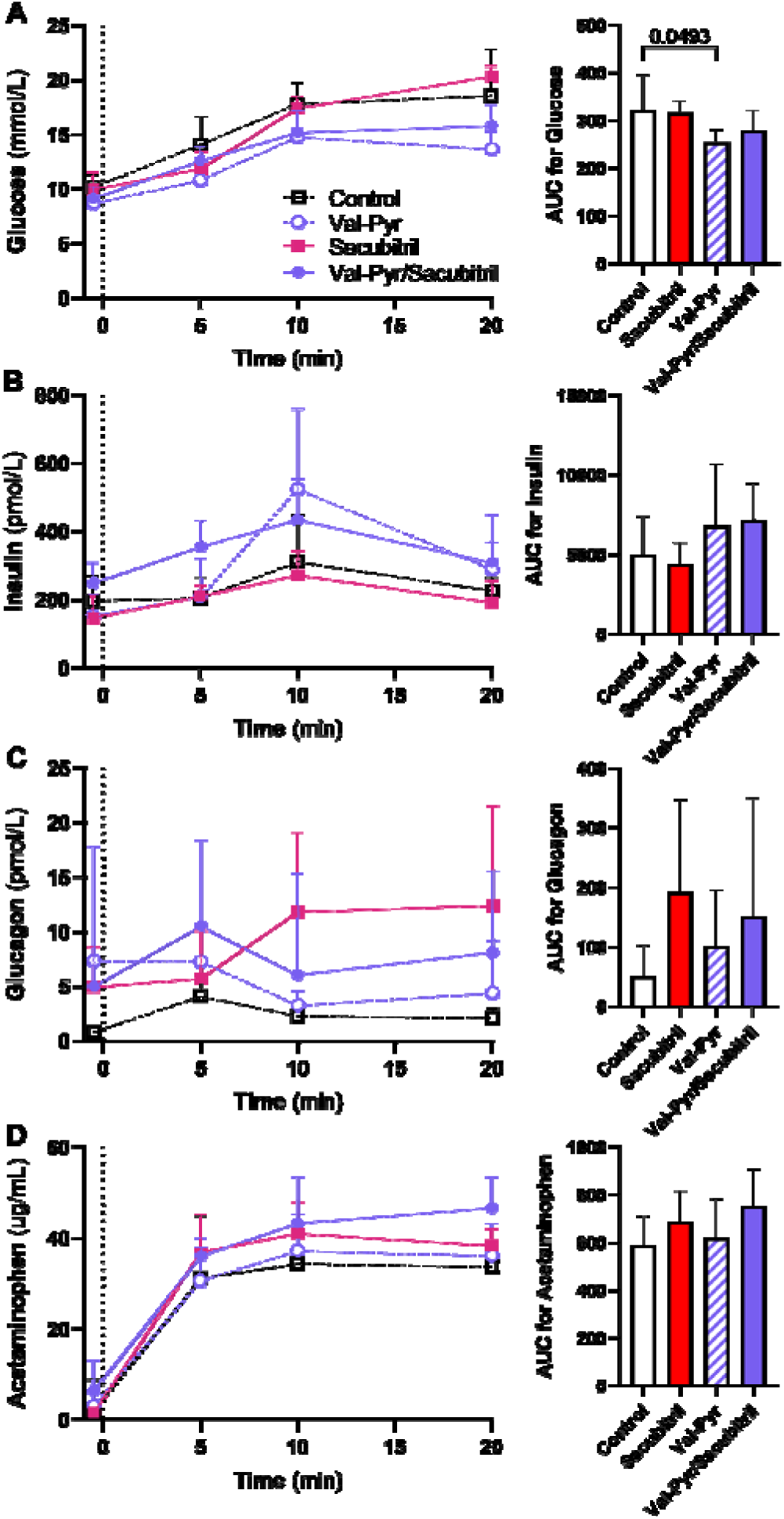
*In vivo* inhibition affects other parameters measured during an OGTT. **Legend:** Effects of *in vivo* inhibition of DPP-4 (Valine pyrrolidide p.o.; purple dashed line, open circle; n=6), NEP (sacubitril p.o.; red solid line, closed square; n=6), both (purple solid line, closed circle; n=6) or control (no inhibitors, black dashed line, open square; n=5) in male C57Bl/6JRj mice on parameters which are frequently assessed with an OGTT. Oral glucose was given at t=0 and blood was drawn from the retrobulbar plexus at t=0, 5, 10 and at 20 min. *Panel (A)* shows the effect on blood glucose levels; *Panel (B)* shows plasma insulin levels; *Panel (C)* shows plasma glucagon levels; and *Panel (D)* shows plasma acetaminophen levels. Acetaminophen was co-administered with the oral glucose load. Values are shown as mean ± SD. Statistical analyses were performed for the AUCs, using one-way ANOVA with post-hoc Šidák focusing only on the difference between inhibitors and control.

### Disappearance of GLP-1(7–36)NH_2_ is strongly reduced by *in vivo* inhibition of DPP-4

To assess the efficacy of *in vivo* inhibition of DPP-4 and NEP, we injected a known amount of GLP-1(7–36)NH_2_ into the tail vein of mice, and measured plasma GLP-1 levels repeatedly afterwards in blood drawn from the retrobulbar plexus (***Figure 6a-b***). Using the Alpco-kit, one minute after injection, GLP-1 levels were significantly higher in sitagliptin versus control (38.6 [18.5 to 58.7] pmol/L; p=0.0002), and versus the combination of sitagliptin/sacubitril (21.0 [0.9 to 41.1] pmol/L, p=0.039) (***Figure 6a***). The subsequent, calculated half-life was 0.89 minutes for control, 1.53 minutes for sitagliptin and 5.13 minutes for sitagliptin/sacubitril. Interestingly, we observed very low levels of total GLP-1 in the same plasma samples when we used the Mercodia-kit (***Figure 6b***). Also, similar to our earlier experiments with the Mercodia-kit, there was no added benefit of DPP-4 inhibition with or without NEP-inhibition.

**Figure 6.**
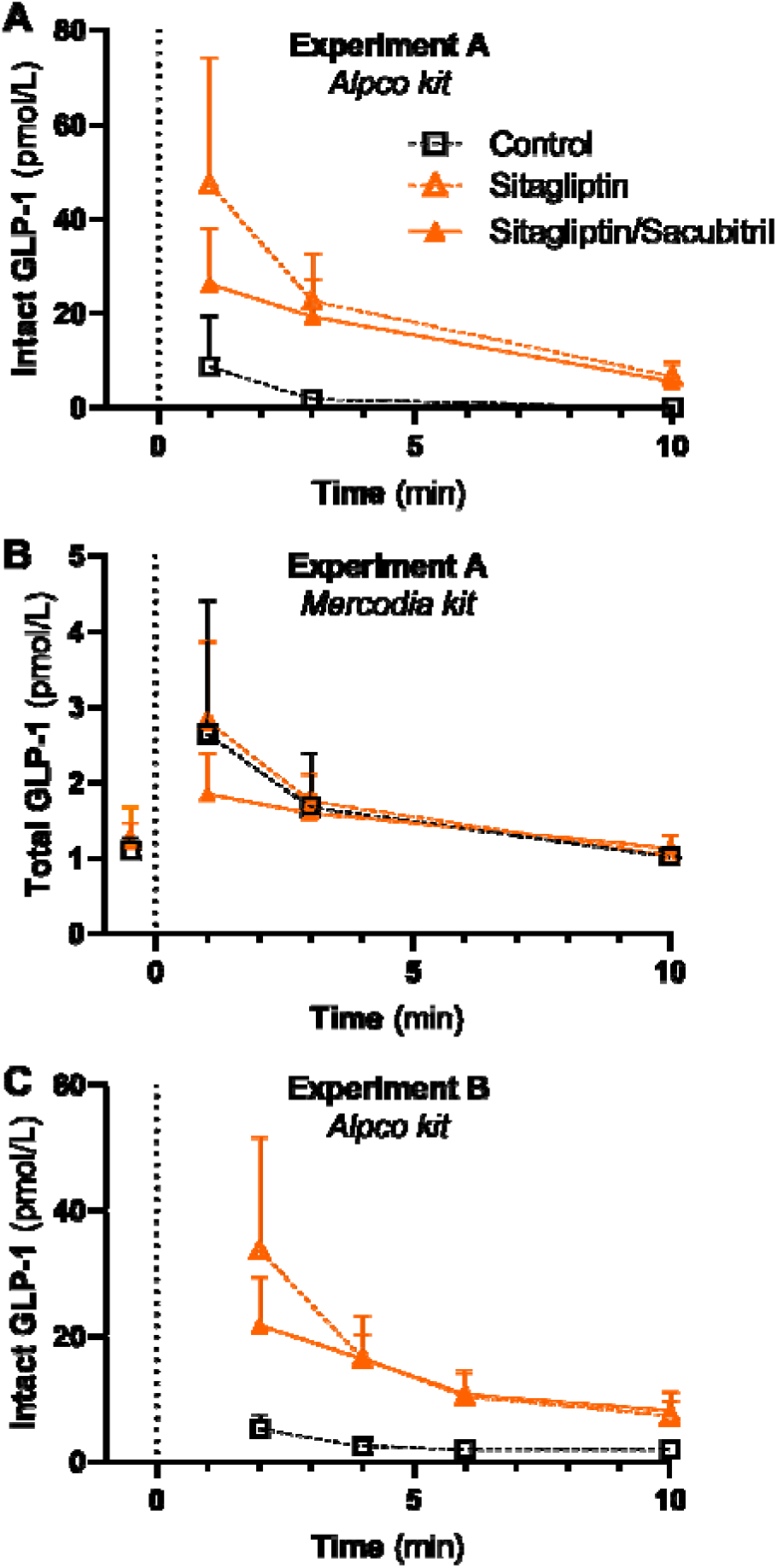
*in vivo* disappearance of GLP-1s. **Legend:** Disappearance of GLP-1 was assessed in two experiments. In *Panel (A)* and *Panel (B)*, 800 fmol of GLP-1(7–36)NH_2_ was injected into the tail vein of male C57Bl6/JRj mice at timepoint t=0, and intact GLP-1 was subsequently measured in plasma using the Alpco kit (panel A) or the Mercodia kit (panel B). This was done without inhibitors (control, black line; n=5), and with DPP-4 inhibition (sitagliptin, orange dashed line; n=5) and DPP-4 and NEP inhibition (sitagliptin/sacubitril, orange solid line; n=5). The expected peak plasma level was ∼300 pmol/L. In *Panel (C)*, a similar experiment was performed, but here GLP-1(7–36)NH_2_ was injected into the vena cava of male C57Bl6/JRj mice (n=4 per group). Values are shown as mean ± SD.

Because of the somewhat unexpected result with a better initial yield with DPP-4 inhibition only versus the cocktail with sacubitril, we performed a second experiment, this time administering GLP-1 directly into the vena cava (ensuring that no technical errors with tail vein injection could occur) (***Figure 6c***). Again, the initial peak was highest in sitagliptin only (reaching a peak value of ∼ 50 pmol/L), followed by sitagliptin/sacubitril. The half-life was 0.79 minutes for control, 1.35 minutes for sitagliptin and 2.63 minutes for sitagliptin/sacubitril.

### *In vitro* degradation of GLP-1

We additionally assessed the *in vitro* degradation of GLP-1(7–36)NH_2_ from mouse plasma, without addition of inhibitors (***Figure 7a***). After administration of GLP-1(7–36)NH_2_, calculated to reach 200 pmol/L, a half-life of 53 min were observed when assessed using the Alpco kit. Interestingly, although we expected similar or even higher levels with the Mercodia kit, as it should measure both GLP-1(7–36)NH_2_ and GLP-1(9–36)NH_2_, the levels of total GLP-1 measured with that kit were ∼100 pmol/L and stable. We next repeated this experiment, but now added GLP-1(9–36)NH_2_ to the mouse plasma (***Figure 7b***). Again, the Mercodia kit did not measure the predicted total GLP-1 level, and levels were stable. As reference, we also used a kit commercially available through IBL, which specifically targets the N-terminus of GLP-1[9-36]NH_2_ and GLP-1[9-37]. GLP-1 levels were slightly higher when measured with the IBL kit, yet the general pattern resembled that as obtained with the Mercodia kit.

**Figure 7.**
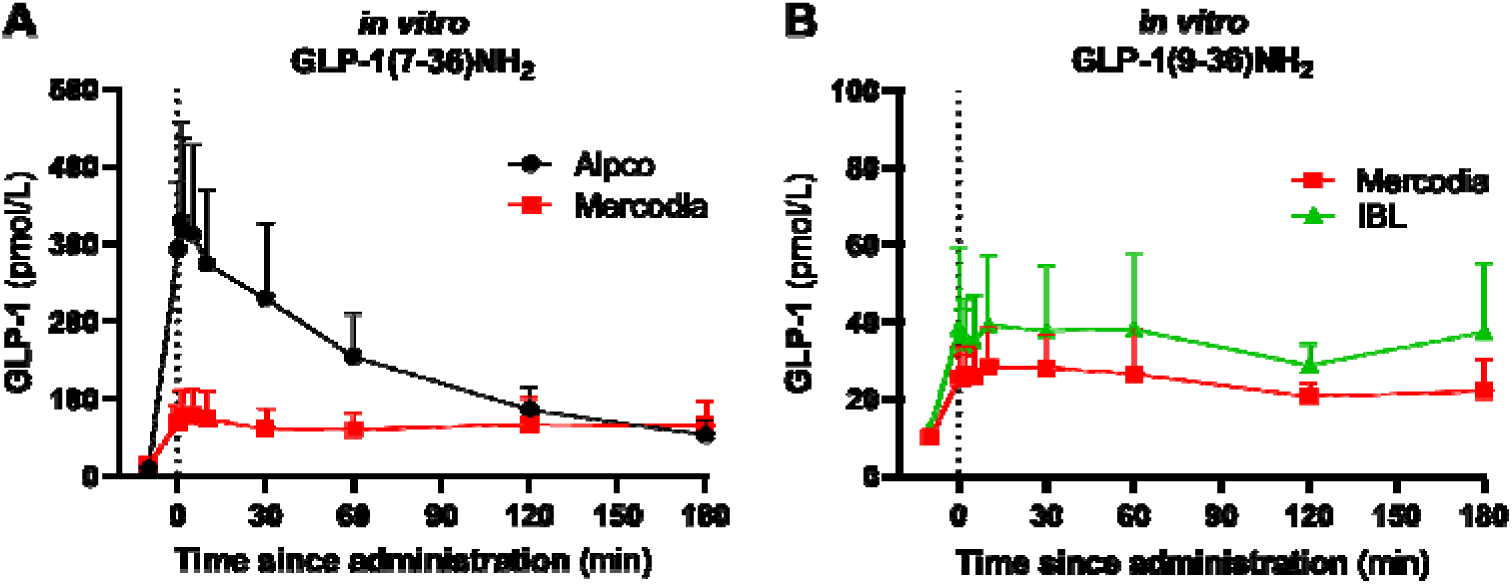
*In vitro* degradation of GLP-1. **Legend:** in *Panel (A)*, GLP-1(7–36)NH_2_ was administered ex vivo to mouse plasma. Plasma was pooled from 5-6 male C57Bl6/JRj mice and divided over 4 tubes to perform these experiment. GLP-1 levels were measured using the Alpco kit (black line) and Mercodia kit (red line) in the same samples, taken before and repeatedly after administration of GLP-1. The expected peak GLP-1 level was 200 pmol/L. in *Panel (B)*, the same experimental setup was used, but here GLP-1(9–36)NH_2_ was administered. GLP-1 was measured using the Mercodia kit, which measures total GLP-1, and an IBL-kit that specifically measures GLP-1(9–36)NH_2_. Values are shown as mean ± SD.

## Discussion / considerations

In the current study, we focused on approaches to estimate glucose stimulated secretion of GLP-1 in mice. Measurements of total GLP-1 levels showed only rapidly occurring (<15 minutes) and small increases after an OGTT. For the measurement of intact GLP-1 levels, *in vivo* inhibition of at least DPP-4, but preferably DPP-4 and NEP appears needed.

Our group has previously shown that it is difficult to measure both intact and total GLP-1 in mice, and that commercially available GLP-1 ELISA kits might not measure levels of this hormone accurately (3). We hypothesized this to be caused by rapid and extensive degradation mediated by both DPP-4 (at the N-terminus side) and NEP (endoproteolytic degradation) (3), leading to several small fragments of the GLP-1 peptide. Especially with sandwich ELISA kits, which rely on one antibody at the C- or N-terminus side, and another antibody of which the region specificity is frequently unknown, peaks of GLP-1 might be missed. This results in low or unmeasurable levels of GLP-1 where one otherwise would have expected to find a peak (3, 5). Importantly, as NEP degrades the GLP-1(9–36) fragment, we anticipated assay difficulties for both intact and total GLP-1.

In the present studies we found only early and very small glucose-induced increases in total GLP-1 levels. Furthermore, attempts to increase the early peak of GLP-1 secretion with *in vivo* NEP inhibition were unsuccessful. When interpreting these data two things must be considered. First, even though an increase in GLP-1 was observed with the Mercodia kit, the absolute levels were low. Moreover, in the *in vitro* degradation studies, the Mercodia kit did not seem to be able to measure GLP-1 quantitatively (expected peak was 200 pmol/L, whereas the measured peak was <80 pmol/L). Since there was a measurable rise in GLP-1 levels after administration of either GLP-1(7–36)NH_2_ or GLP-1(9–36)NH_2_, the Mercodia kit apparently picks up something, yet not to the same extent as other kits (as seen with the Alpco and IBL kits). Second, an important consideration when using a total GLP-1 kit and early time-points, is that the assay also picks up N-terminally extended GLP-1(1–36)NH_2_. This extended and inactive form of GLP-1 is co-produced in the pancreas with glucagon (12), and is likely responsible for the fast peak (within 5 minutes) seen after a stressful stimulus (for example due to oral gavage and blood sampling) (3). GLP-1(1–36)NH_2_ does not affect insulin secretion (13), and is not the form of GLP-1 most researchers are interested in when studying ‘GLP-1 secretion’. Whether such stress-induced GLP-1(1–36)NH_2_ disqualifies the total GLP-1 measurement was not the topic of this study, but we advise authors to at least think about this possibility when using such kits.

The lack of response of intact GLP-1 during glucose stimulation is probably due to the susceptibility to degradation by DPP-4 and NEP (3, 14). It did not increase the levels of measurable GLP-1 when the blood samples were immerged in tubes containing protease-inhibitors (‘ex vivo inhibition’). This is underscored by the strong discrepancy we found between *in vivo* disappearance and *in vitro* plasma disappearance of GLP-1(7–36)NH_2_, which suggests that disappearance occurs rapidly *in vivo* by processes not restricted to plasma. Therefore, inhibiting enzymes in plasma *after* blood withdrawal is not sufficient to stabilize GLP-1.

In contrast, by inhibiting DPP-4 and NEP *in vivo*, the degradation of secreted intact GLP-1 is immediately reduced, allowing measurement by sandwich ELISA. The potency of *in vivo* inhibition was further demonstrated by injecting a known amount of GLP-1(7–36)NH_2_, and assessing subsequent disappearance. While the post-injection peak was comparable between the control group and the inhibitor groups, the degradation in the control group was profound. Inhibition of DPP-4, with or without additional inhibition of NEP, led to stable levels of intact GLP-1 (at least during the 10 minute observation period). Importantly, while we have shown that the *in vivo* inhibition technique with valine pyrrolidide/sacubitril and sitagliptin/sacubitril works well with the Alpco kit, this does not guarantee that the same is true when using other combinations.

The experiments with injection of exogenous GLP-1(7–36)NH_2_ *in vivo* are important as they address the question whether it is at all possible to measure a release of endogenous GLP-1 to the circulation. The experiments confirmed the very high disappearance rate of GLP-1 (T½ ∼ 2 min) but also that any GLP-1 that reaches the circulation should be measurable after addition of the inhibitors.

According to the manufacturer, the Alpco kit only measures GLP-1(7–36)NH_2_, with no cross-reactivity to GLP-1(9–36)NH_2_ or the glycine-extended GLP-1(7–37). Although we did not confirm this in our study, this information suggests that the combination of antibodies targets both the N-terminus and C-terminus of GLP-1 (as a combination of antibodies directed at the N-terminus and the mid-region would also measure GLP-1[7-37]). From a methodological point of view this is interesting, as it shows that our *in vivo* inhibition technique stabilizes the entire GLP-1(7–36)NH_2_ peptide.

An important point of discussion is how to now interpret the vast body of literature where GLP-1 is measured in mice without the here introduced approaches. As stated before, levels could be false-negative, false-positive, or accurate. Therefore, we suggest that readers carefully read the methodology of the paper and assess the timing and height of the GLP-1 peak (absolute values) and the alignment with glucose (or whatever stimulus is employed) and insulin peaks. If GLP-1 levels are expressed relative to baseline, this may indicate that the levels were not in the physiological range. Yet a large portion of papers do show absolute levels, in the expected range (that is: comparable to humans). This can only be explained if the applied ELISA kit is able to measure GLP-1 fragments, and as such does not require *in vivo* inhibition. As over 30 commercial kits are available, we did not assess the efficacy of each kit. It is also crucial to emphasize that we only validated the *in vivo* inhibition technique with valine pyrrolidide/sacubitril and sitagliptin/sacubitril with the Alpco kit. We strongly urge others who want to use different inhibitors or different ELISA kits to first perform a pilot study and assess the resulting GLP-1 response. Finally, we only used male C57Bl/6JRj mice, and thus cannot say whether this method works in female animals and other strains.

*In vivo* inhibition of DPP-4 and NEP also affects associated physiological processes. Indeed, although the numbers were small, precluding robust statistical analysis, we observed the expected increase in insulin with DPP-4 inhibition, and lower levels of glucagon and glucose (8, 9). In contrast, inhibition of NEP increased glucagon and glucose levels, as also seen previously (10, 11). The lowering effect of DPP-4 inhibition on glucagon levels was presumably counteracted by the stabilization provided by NEP-inhibition. As a result, with the inhibitor cocktail, glucose levels after OGTT were comparable to the control group. Finally, the combinations with sacubitril appear to augment gastric emptying rate, as acetaminophen levels rose slightly faster in those groups.

In conclusion, we here demonstrated approaches to measure plasma levels of total and intact GLP-1 during an OGTT in male C57Bl/6JRj mice with commercially available ELISA kits. Our suggestion is to measure intact GLP-1 (GLP-1[7-36]NH_2_) during *in vivo* inhibition of DPP-4 (and perhaps also NEP) thus stabilizing the GLP-1 peptide. Measuring intact GLP-1, and not total GLP-1, diminishes the risk of measuring GLP-1(1–36)NH_2_ (depending on the assay). While *in vivo* inhibition affects other parameters normally tested during an OGTT (insulin and glucagon), it allows smaller plasma volumes due to assessment using ELISA kits, and thus reduced animal burden. This approach will aid future studies of GLP-1 secretion in mice.

## Grants

This study was funded by ERC and Novo Nordisk Foundation. MMS is a Lundbeck postdoc fellowship recipient.

## Disclosures

MMS, KDG, BH and SLJ: Nothing to disclose

JJH: Advisory boards: Novo Nordisk; Lecture fees: Novo Nordisk; Merck

NJWA: Advisory boards: Novo Nordisk; Boehringer Ingelheim, Lectures Fees: Novo Nordisk, Merck, Mercodia, Research Support: Novo Nordisk, MSD, Mercodia, Roche, EvoSep

## Acknowledgements

None

## Funding / Conflicts of interest

See title page

---

**Supplemental Figure 1.**
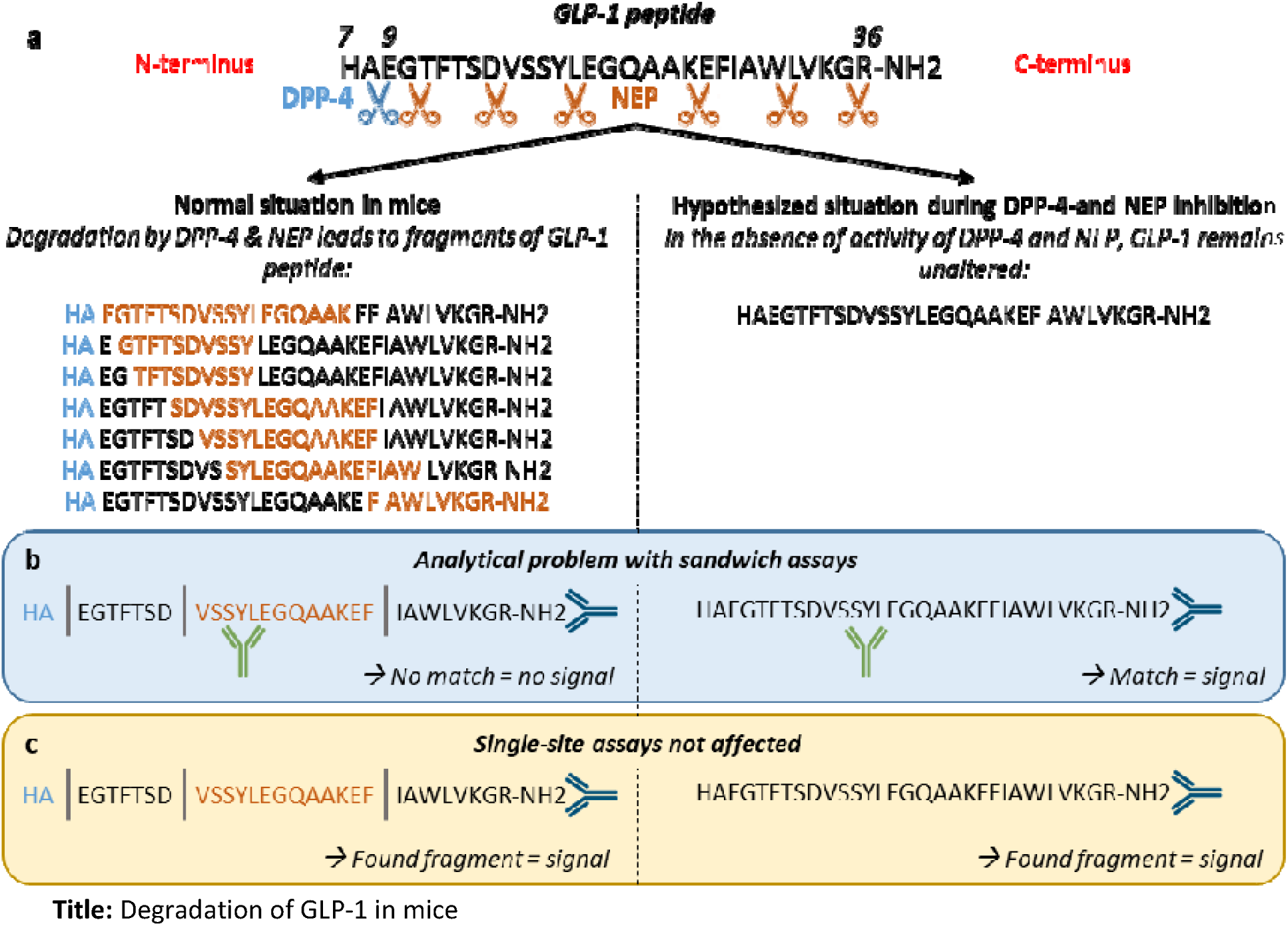
Degradation of GLP-1 in mice. **Legend:** Panel (a): In mice, two separate and simultaneous processes rapidly degrade GLP-1. First, DPP-4 removes histidine (H) and alanine (A) from the N-terminal part of GLP-1(7–36)NH_2_, creating GLP-1(9–36)NH_2_ (the blue scissor indicates the cleavage site). Simultaneously, neprilysin (NEP) activity leads to endoproteolytic degradation of GLP-1, creating several smaller fragments (the red scissors indicate the cleavage points; only the most common fragments are depicted here in red) (3)). Hypothetically, adding inhibitors of DPP-4 and NEP will reduce this degradation, leaving only the intact GLP-1 peptide (shown on the right). Panel (b): The rapid and extensive degradation of GLP-1 creates an analytical problem with sandwich assays, which require that two antibodies target the same fragment of the peptide. If the antibodies target enzymatically separated fragments, the assay will not detect GLP-1. We hypothesized that stabilizing the GLP-1 peptide using specific DPP-4 and NEP inhibitors would allow adequate measurements using sandwich assays. The “specific” assays are supposed to have one terminal wrapping antibody although it may in fact be “sideviewing” (but near the terminal); this information is not provided by the manufacturer. The other antibody is probably mainly sideviewing, but again exactly which part of the molecule is bound is frequently unknown or not disclosed. Depicted is an example of one fragment generated by NEP cleavage, where the vertical lines indicate the NEP cleavage points. Panel (c): Single-site assays rely on only one antibody to measure GLP-1, and are not disturbed by the fragmentation of the peptide. However, assays based on this technique currently require a large plasma volume and lack sensitivity. Figure based on (3).

**Supplemental Figure 2.**
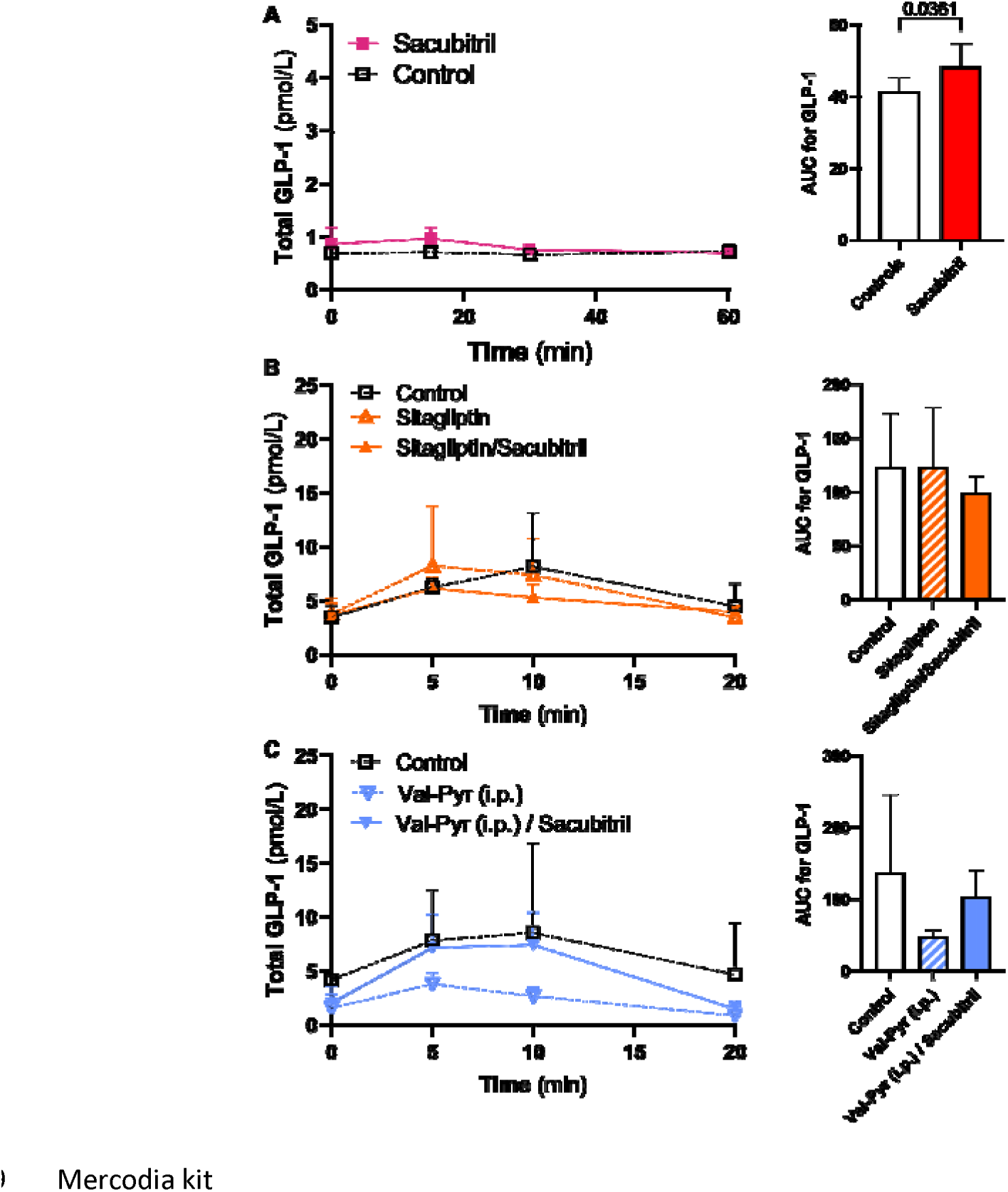
*In vivo* enzyme inhibition does not lead to better results when measuring total GLP-1 with the Mercodia kit. **Legend:** Effects of *in vivo* inhibition of DPP-4 (Valine pyrrolidide p.o.) and/or NEP (sacubitril) on total GLP-1 levels as measured with the Mercodia assay in male C57Bl/6JRj mice. Panel (A) shows the effect of NEP-inhibition alone (sacubitril; red solid line, closed squares; n=6) versus control (no inhibition, black dashed line, open squares; n=7); Panel (B) shows the effect with sitagliptin (orange dashed line, open triangle; n=5) and sitagliptin/sacubitril (orange solid line, closed triangles; n=5) and control (n=5); and *Panel (C)* shows the effect of Valine pyrrolidide i.p. (blue dashed line, open triangles; n=5), Valine pyrrolidide/sacubitril (blue solid line, closed triangles, n=5) and control (n=3). Values are shown as mean ± SD. Statistics were performed using an unpaired t-test (comparing 2 groups) or a one-way ANOVA (comparing 3 groups), with post-hoc Šidák focusing only on the difference between inhibitors and control.

